## Supplementary material for "Balancing central control and sensory feedback produces adaptable and robust locomotor patterns in a spiking, neuromechanical model of the salamander spinal cord": S1_signal_processing

### Neural metrics

In each model simulation, several metrics were computed to evaluate the performance of the spiking neural network. A schematic of the signal processing pipeline is drawn in Fig S1A. Spikes are counted at each time-step and for each equivalent hemisegment to obtain the spike count signals $SC_i$ (see Fig S1C, middle plot). Then, a cubic spline fit [1] is applied to this data in order to obtain a filtered spike count signal for each hemisegment $FSC_i$ (see Fig S1C, bottom plot). The frequency of the oscillations ($f_{neur}$) is computed from the location of the maximum in the amplitude diagram of the Fourier Transform ($FT$) of the corresponding signals, as

$$f_{neur} = \frac{1}{2\pi} < \arg\max_{\omega}(|FSC_i(\omega)|) >_i \tag{1}$$

Where $< \cdot >_i$ denotes the average computed across all the segments. The intersegmental phase lag (IPL) between segments is computed based on the location of the maximum of the cross-correlation between the $SC$ signals of adjacent segments as

$$IPL = f_{neur} < \arg\max_{t}(xcorr(FSC_i, FSC_{i+1})(t)) >_i \tag{2}$$

Note that phase lags are normalized by the frequency of the neural oscillations. Similarly, the total wave lag of the neural signal ($TWL$) is computed as the sum of the IPL across the network

$$TWL = f_{neur} \sum_i \arg\max_{t}(xcorr(FSC_i, FSC_{i+1})(t)) \tag{3}$$

This quantity (often referred to as wave number) represents the number of travelling waves present along the axial network at the any given time. A total wave lag of 0 corresponds to a standing wave, resulting in a synchronous activation of all muscles on one side of the body. A total wave lag of 1 indicates that, in each cycle, the body generates a complete wave with a 100% phase lag between head and tail segments. the activation of the head segment will occur immediately after the inactivation of the last tail segment. The peak-to-through correlation coefficient (PTCC) is computed as the difference between the first two extrema of the autocorrelogram of the corresponding segmental signal.

$$PTCC =< \max_{t}(corr(FSC_i)(t)) - \min_{t}(corr(FSC_i)(t)) >_i \tag{4}$$

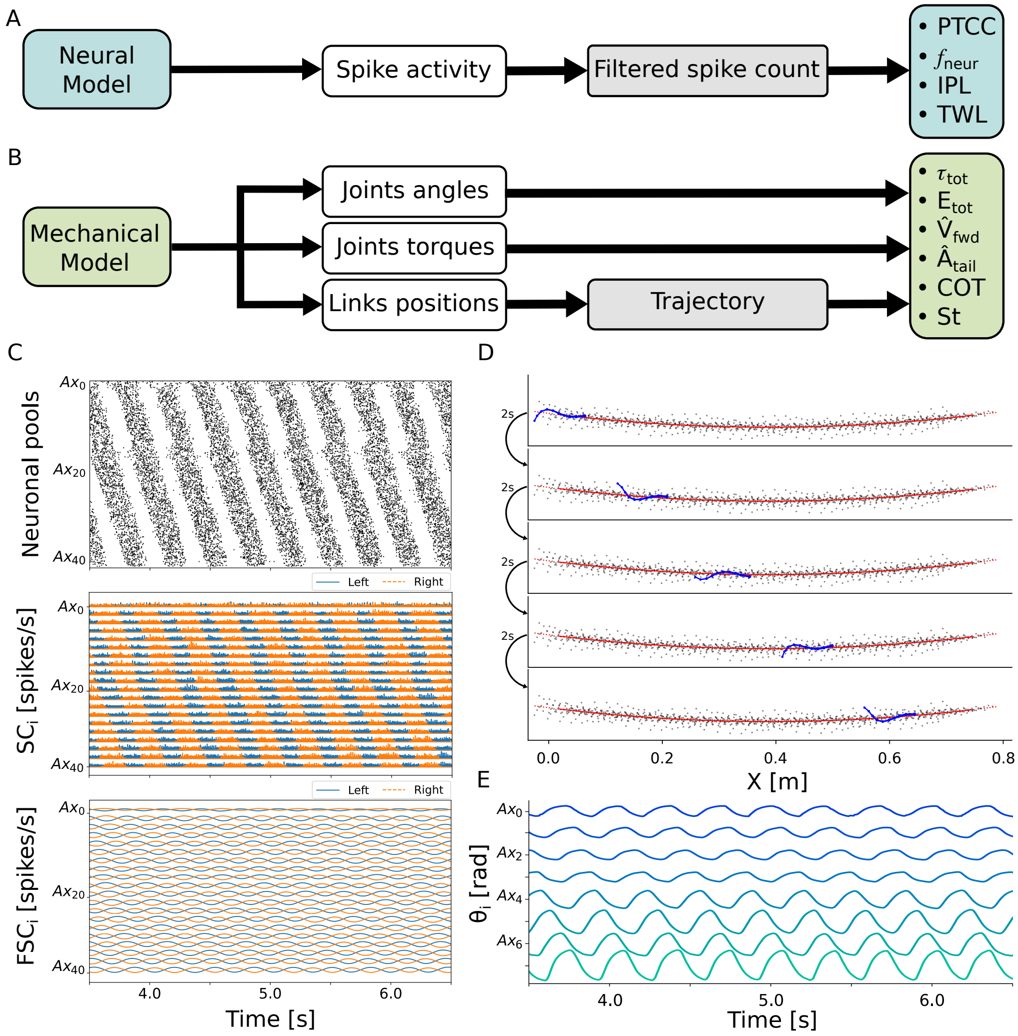


**Fig. S1. Signal processing pipeline. A**) shows the pipeline for the computation of the neural metrics. The discrete spike activity of the network is converted into smooth signals (filtered spike counts) for further processing. **B**) shows the pipeline for the computation of the mechanical metrics. The trajectory line is computed based on the evolution of the links positions. The trajectory is combined with the joint angles and torques evolution to compute the metrics. **C**) shows the discrete firing activity of the CPG network (top figure), the evolution of the spike count signal for each equivalent segment (middle figure) and the evolution of the filtered spike counts (bottom figure). **D**) displays the trajectory line (in red) fitting the coordinates of all the links during the simulation (in grey). The snapshots show the body configuration at different times of the simulation, with an interval of 2s between each frame. **E**) shows the evolution of the joint angles during the simulation. Note that the caudal joints display larger angular excursions with respect to the rostral joints. This is a typical feature of anguilliform swimming kinematics.

A PTCC value of 2 corresponds to a perfectly periodic signal, while lower PTCC values correspond to increasingly less periodic signals. For this reason, PTCC is used to evaluate the degree of periodicity of the segmental activations during the experiments [2]. Higher periodicity is interpreted as a higher stability of the corresponding patterns.

### Mechanical metrics

Similarly to the neural metrics, several quantities were computed to evaluate the performance of the mechanical model. A schematics of the signal processing pipeline is drawn in Fig S1B. For each simulation, the trajectory curve denotes the parabolic line fitting the coordinates of all the body links (see Fig S1D). The trajectory curve is computed as

$$\{a_2^*, a_1^*, a_0^*\} = \arg\min_{a_2,a_1,a_0} (MSE(a_2, a_1, a_0)) \tag{5}$$

Where the curve is expressed by the quadratic function

$$y = a_2 x^2 + a_1 x + a_0 \tag{6}$$

And MSE is the mean square error between the link coordinates and the quadratic curve

$$MSE(a_2, a_1, a_0) = \sum_t \sum_i (x_i(t) - x)^2 + (y_i(t) - y)^2 \tag{7}$$

The evolution of the lateral displacement of every link is computed as the distance between the link coordinates and the trajectory line.

$$d_i(t) = \left| \begin{pmatrix} x_i(t) \\ y_i(t) \end{pmatrix} - \arg\min_{x,y} \left| \begin{matrix} x_i(t) - x \\ y_i(t) - a_2^* x^2 - a_1^* x - a_0^* \end{matrix} \right| \right| \tag{8}$$

The amplitudes of such displacements are computed from the maximum of the Fourier Transform of each signal as

$$A_i = \max_\omega (|d_i(\omega)|) \tag{9}$$

Where the value corresponding to the last joint represents the tail beat amplitude $A_{tail}$.

The amplitude of each joint oscillation angle ($\Theta_i$) is computed analogously from the Fourier transform of its temporal evolution (see Fig S1E) as

$$\Theta_i = \max_\omega (|\theta_i(\omega)|) \tag{10}$$

The center of mass position $\mathbf{X}_{COM}(t)$ is computed as the average of the body link positions $\mathbf{X}_i(t)$ at time $t$ weighted by the corresponding links masses $m_i(t)$ according to

$$\mathbf{X}_{COM}(t) = \begin{pmatrix} x_{com}(t) \\ y_{com}(t) \end{pmatrix} = \frac{\sum_i m_i \mathbf{X}_i(t)}{\sum_i m_i} \tag{11}$$

The forward trajectory direction $\mathbf{w}_{fwd}$ is computed from the first principal component of the $2X9N$ matrix $\mathbf{X}$ containing the coordinates of the 9 axial link across the N steps of the simulation, according to

$$\mathbf{w}_{fwd} = arg \max_{\|\mathbf{w}\|=1} \{\mathbf{w}^T \mathbf{X}^T \mathbf{X} \mathbf{w}\} \tag{12}$$

The forward speed $V_{fwd}$ of the mechanical model is then obtained from the projection of $\mathbf{X}_{COM}$ along the trajectory line according to

$$V_{fwd} = \mathbf{w}_{fwd}^T \frac{\mathbf{X}_{COM}(N\Delta t) - \mathbf{X}_{COM}(0)}{N\Delta t} \tag{13}$$

Where $\Delta t$ is the timestep of the mechanical simulation.

The mechanical metrics listed so far are also normalized by the length $L$ of the simulated body (10 cm) for a better comparison with biological experiments. This corresponds to the computation of specific forward speed $\hat{V}_{fwd}$ and specific tail beat amplitude $\hat{A}_{tail}$, defined as

$$\hat{V}_{fwd} = V_{fwd}/L \tag{14}$$

$$\hat{A}_{tail} = A_{tail}/L \tag{15}$$

The Strouhal number ($St$) is computed as

$$St = 2fA_{tail}/V_{fwd} \tag{16}$$

It represents a non-dimensional quantity that is widely used to evaluate the swimming performance of animals. Most studies report that animals tend to swim with Strouhal numbers in the range 0.2 to 0.4 [3]. Salamanders, on the other hand, appear to express higher St values (0.516 in [4], 0.578 in [5]), possibly denoting a lower swimming efficiency.

The total torque consumption ($\tau_{tot}$) is obtained by integrating the absolute values of the active torques exerted by the joints across the entire duration of the simulation

$$\tau_{tot} = \sum_t \sum_i \alpha_i |M_{diff_i}(t)| \tag{17}$$

It is used as an indication of the amount of effort required to produce the locomotor patterns. The total energy consumption ($E_{tot}$) is computed from the integral across the duration of the simulation of the product between the active torque exerted by the joints and their instantaneous rotational speed.

$$E_{tot} = \sum_t \sum_i \alpha_i M_{diff_i}(t) \dot{\theta}_i(t) \tag{18}$$

The Cost of Transport ($COT$) is computed as the ratio between the total energy consumption and the forward speed of the mechanical model.

$$COT = E_{tot}/V_{fwd} \tag{19}$$

The cost of transport represents a metrics for the energy efficiency of the generated locomotion pattern. More efficient locomotor patterns will produce faster locomotor speeds with lower energy consumptions, thus resulting in lower COT values.
