## Supplementary material for "Balancing central control and sensory feedback produces adaptable and robust locomotor patterns in a spiking, neuromechanical model of the salamander spinal cord": S2_neuronal_synaptic_properties

### Neuronal and synaptic parameters

#### Neuronal populations

The network comprises 5 different classes of neurons, namely excitatory neurons (EN), inhibitory neurons (IN), motoneurons (MN), reticulospinal neurons (RS) and propriosensory neurons (PS). The neuronal classes differ for their neuronal and synaptic properties, which were obtained based on experimental data and previous simulation studies (see Table S1 and Table S2). Unknown properties were treated as open parameters to manually tune the activation of the network according to the known salamander locomotor patterns.

**Table S1. Neuronal Parameters**

| Neuron Type | $t^*_{\text{refr}}$ (ms) | $\tau^*_{\text{memb}}$ (ms) | $R^*_{\text{memb}}$ (G$\Omega$) | $\Delta\omega$ (pA) | $\tau_\omega$ (pA) |
|---|---|---|---|---|---|
| $EN_{axial}$ [1–3] | 5.0 | 26.8 | 1.6 | 3.0 | 200.0 |
| $EN_{limbs}$ [1–3] | 5.0 | 37.5 | 2.1 | 0.7 | 500.0 |
| $IN_{axial}$ [1–3] | 5.0 | 26.8 | 1.6 | 3.0 | 200.0 |
| $IN_{limbs}$ [1–3] | 5.0 | 37.5 | 2.1 | 0.7 | 500.0 |
| RS [4] | 5.0 | 26.8 | 8.0 | 0.0 | - |
| MN [5] | 5.0 | 26.8 | 16.0 | 0.0 | - |
| PS [6, 7] | 50.0 | 26.8 | 0.6 | 0.0 | - |

In Table S1, the value of the variables denoted by an asterisk was sampled randomly from a Gaussian distribution for each neuron of the network. The table reports the mean of the distributions. The standard deviation was set to 20% of mean value. The value of $E_{rest}$, $E_{reset}$, $E_{thres}$ (see Eq 1 in the main text) was shared among all the neuron populations and equal to $-70.0mV$, $-70.0mV$ and $-38.0mV$ respectively. The RS, MN and PS populations were modeled with non-adaptive neurons $\Delta\omega = 0$. The PS population was modeled with a large $t_{refr}$ value to limit its maximum firing rate according to experimental findings [6, 7]. The same effect could be obtained using highly adaptable neurons and a more physiologically-sound refractory period.

In Table S1, the parameters for the synaptic variables of the model are reported (see Eq 3,4 in the main text). Neurons with an excitatory action acted on the AMPA and NMDA synaptic conductances. Conversely, inhibitory neurons influenced the GLYC component.

**Table S2. Synaptic Parameters**

| Synaptic Type | $E_{\text{syn}}(mV)$ | $\tau_{\text{syn}}(ms)$ | Neuron populations |
|---|---|---|---|
| AMPA [1, 2] | 0.0 | 20.0 | EN, RS, PS |
| NMDA [1, 2] | 0.0 | 100.0 | EN, RS, PS |
| GLYC [1, 2] | -85.0 | 20.0 | IN, PS |

Finally, the synaptic weights $\Delta g_{syn}$ (see Eq 4 in the main text) were selected to replicate the patterns of activations observed in salamanders and the values are reported in Table S3.

**Table S3. Synaptic Weights**

| Source | Target | $\Delta g_{ampa}$ | $\Delta g_{nmda}$ | $\Delta g_{glyc}$ |
|---|---|---|---|---|
| $EN_{axial}$ | $CPG_{axial}$ | 0.025 | 0.007 | - |
| $IN_{axial}$ | $CPG_{axial}$ | - | - | 0.036 |
| $EN_{limbs}$ | $CPG_{limbs}$ | 0.083 | 0.025 | - |
| $IN_{limbs}$ | $CPG_{limbs}$ | - | - | 0.128 |
| $EN_{limbs}$ | $CPG_{axial}$ | 0.044 | 0.013 | - |
| $IN_{limbs}$ | $CPG_{axial}$ | - | - | 0.086 |
| $RS_{axial}$ | $CPG_{axial}$ | 0.053 | 0.015 | - |
| $RS_{limbs}$ | $CPG_{limbs}$ | 0.083 | 0.025 | - |
| $PS^*_{EX}$ | CPG | 0.025 $\omega_{PS}$ | 0.007 $\omega_{PS}$ | - |
| $PS^*_{EX}$ | MN | 0.025 $\omega_{PS}$ | 0.007 $\omega_{PS}$ | - |
| $PS^*_{IN}$ | CPG | - | - | 0.036 $\omega_{PS}$ |
| $PS^*_{IN}$ | MN | - | - | 0.036 $\omega_{PS}$ |
| EN | MN | 0.070 | 0.020 | - |
| MN | MC | 0.350 | 0.150 | - |

Despite having identical neuronal properties, the IN and EN populations differ in their synaptic actions (see Table S2). Together, the two populations represent the core rhythmogenic component of the network (i.e., the CPG network). For this reason, the EN and IN populations were assigned different values for their axial and limbs sub-networks [1, 8]. The primary distinguishing factor between the two sub-populations lies in their adaptation time-constants and weights (see Table S1 and Table S3). The higher adaptation time of the limb populations results in a lower frequency of the generated oscillations, in accordance with experimental findings.

Note that the synaptic weight of excitatory ($PS_{EX}$) and inhibitory ($PS_{IN}$) proprioceptive sensory neurons is scaled by the sensory feedback weight $\omega_{PS}$. When $\omega_{PS} = 1$, the sensory neurons have the same synaptic weight as the intra-CPG connections between EX and IN neurons.

#### Muscle cells model

The muscle cells act as low-pass filters of the input activity (see Eq 6 in the main text). Their input is normalized in the range [0, 1] and can be interpreted as the degree of activation of the corresponding network hemisegment. The parameters for the muscle cells are reported in Table S4.

**Table S4. Muscle cells Parameters**

| Parameters | Value |
|---|---|
| $\tau_{\mathrm{mc}}(ms)$ | 100.0 |
| $E_{\mathrm{AMPA}_{mc}}(\#)$ | 1.0 |
| $E_{\mathrm{NMDA}_{mc}}(\#)$ | 1.0 |
| $E_{\mathrm{GLYC}_{mc}}(\#)$ | 0.0 |
| $\tau_{\mathrm{AMPA}_{mc}}(ms)$ | 2.0 |
| $\tau_{\mathrm{NMDA}_{mc}}(ms)$ | 10.0 |
| $\tau_{\mathrm{GLYC}_{mc}}(ms)$ | 2.0 |

**Connectivity parameters**

**Table S5. RS connectivity**

| RS population | Amp | $UP_0$ [m] | $UP_1$ [m] | $DW_0$ [m] | $DW_1$ [m] |
|---|---|---|---|---|---|
| Rost | 0.02 | - | 0.000 | 0.125 | 0.225 |
| Midt | 0.02 | 0.025 | 0.125 | 0.250 | 0.350 |
| Endt | 0.02 | 0.150 | 0.250 | 0.375 | 0.400 |
| Pelv | 0.02 | 0.350 | 0.375 | 0.583 | 0.683 |
| Tail | 0.02 | 0.483 | 0.583 | 0.792 | 0.892 |
| Caud | 0.02 | 0.692 | 0.792 | 1.000 | - |

The RS connection probabilities are given by overlapping trapezoidal distributions. The probability of a connection between neurons increases linearly from 0 to the maximum amplitude (Amp) for neurons placed in a position between $UP_0$ and $UP_1$. The probability of connections is constant and equal to Amp for connections between $UP_1$ and $DW_0$. Finally, the connection probability decreases linearly from Amp to 0 for neurons in positions between $DW_0$ and $DW_1$.

**Table S6. Axial connectivity**

| Source | Target | Side | A | $\sigma_{up}$ (mm) | $\sigma_{dw}$ (mm) |
|---|---|---|---|---|---|
| EN | EN | IPSI | 0.5 | 2.00 | 3.00 |
| EN | IN | IPSI | 0.5 | 2.00 | 2.50 |
| IN | EN, IN | CONTRA | 0.5 | 2.00 | 3.00 |
| EN | MN | IPSI | 0.5 | 2.50 | 5.00 |
| IN | MN | CONTRA | 0.5 | 1.25 | 2.50 |
| MN | MC | IPSI | 1.0 | 2.50 | 2.50 |
| $PS$ | EN, IN, MN | IPSI, CONTRA | 1.0 | 5.00 | 5.00 |
