## Supplementary material for "Balancing central control and sensory feedback produces adaptable and robust locomotor patterns in a spiking, neuromechanical model of the salamander spinal cord": S3_muscle_properties_optimization

### Optimization of the muscle properties

Muscle properties encode the way the neuronal signals are translated into a mechanical output. For this reason, the selection of biologically adherent muscle parameters is fundamental to ensure the plausibility of the modeled behaviors. Additionally, the joints along the spinal cord are subjected to different inertial forces during locomotion due to their different location along the kinematic chain. The inertial forces can vary several orders of magnitude between the joints located in the trunk and the ones located in the tail. For this reason, choosing constant muscle parameter values across the axial network would negatively affect the plausibility of the results. Indeed, in that case the active and passive torques exerted by the muscles would either be scaled for joints subjected to high inertial forces or to low inertial forces, resulting inadequate for the others.

In this context, the linearity of the Ekeberg muscle model for constant co-contraction values allows to recast the problem of muscle parameters selection into a more tractable one. Different muscle parameters can be selected for different joints so that all the muscles will exert their action in a dynamical regime where passive forces, active forces and inertial forces all play a relevant role [1, 2].

For the optimization, we considered the activation of an individual axial joint at a time (see Supp. Fig S1A). In first approximation, the joint will respond to an input signal as a spring-mass-damper system with spring stiffness K, damping C and mass M. The transfer function of the equivalent linear time-invariant dynamical system can be expressed as

$$H(s) = \frac{\alpha\theta(s)}{M_{diff}(s)} = \frac{\frac{\alpha}{M}}{s^2 + \frac{C}{M}s + \frac{K}{M}} = G_0 \frac{\omega_n^2}{s^2 + 2\zeta\omega_n s + \omega_n^2} \tag{1}$$

Where $\theta(s)$ and $M_{diff}(s)$ are the laplace transforms of the joint angle and joint motor signal flexor-extensor difference, respectively. From the system parameters, we can define the natural frequency $F_n$, damping ratio $\zeta$ and zero frequency gain $G_0$ as

$$F_n = \frac{1}{2\pi}\sqrt{\frac{K}{M}} \tag{2}$$

$$\zeta = \frac{C}{2\sqrt{KM}} \tag{3}$$

$$G_0 = \frac{\alpha}{K} \tag{4}$$

Note that for a constant input signal $M_{diff}$ the system reaches the equilibrium with a deflection equal to $M_{diff}\frac{\alpha}{\beta}$. From this observation, it follows that the desired zero-frequency gain value $G_0^*$ can be obtained by setting

$$\alpha = G_0^* \beta \tag{5}$$

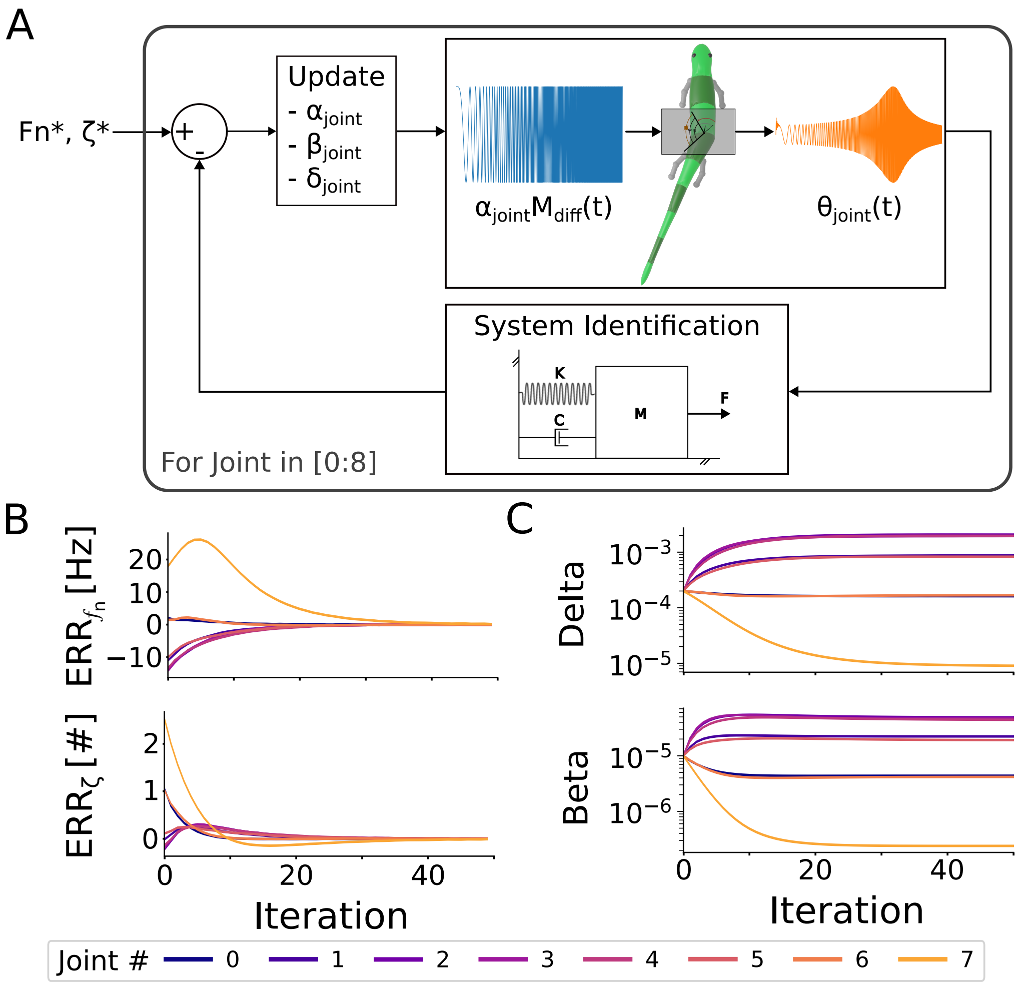


**Fig. S1. Optimization of mechanical model parameters. A**) shows the pipeline for the optimization of the muscle parameters. At each iteration, a sweep activation is given as input to a single joint and the corresponding angle evolution is simulated. The parameters of an equivalent spring-mass-damper system are computed via system identification. Finally, the differences between the estimated and target resonance frequency ($F_n$) and damping ratio ($\zeta$) are used to update the parameters of the muscle model ($\alpha_{joint}$, $\beta_{joint}$, $\delta_{joint}$). **B**) shows the evolution of the error for $F_n$ (top) and $\zeta$ (bottom) during the optimization. It can be noted that the error converges to zero for all the joints and for both parameters. **C**) shows the evolution of the $\delta$ (top) and $\beta$ (bottom) parameters of all the axial joints during the optimization.

Thus, during the optimization, the value of $\alpha$ was scaled according to the modifications made to the value of $\beta$. At every optimization iteration, we stimulated each joint with a sweep activation signal ranging from 0Hz to 5Hz over 30s and recorded the corresponding output angle. Consequently, we performed a parameter identification with Python library sisidentpy [3] in order to compute the parameters of the equivalent spring mass damper system of every joint. From the estimated parameters, we computed the following relative errors

$$ERR_{F_n} = \frac{F_n - F_n^*}{F_n^*} \tag{6}$$

$$ERR_\zeta = \frac{\zeta - \zeta^*}{\zeta^*} \tag{7}$$

Where $F_n^*$, $\zeta^*$ are the target resonance frequency and damping ratio, respectively. Based on the error computed with the equivalent parameters, we modified the parameters of the Ekeberg muscle model of every joint according to

$$\alpha' = \alpha(1 - ERR_{F_n}) \tag{8}$$

$$\beta' = \beta(1 - ERR_{F_n}) \tag{9}$$

$$\zeta' = \zeta(1 - ERR_{F_n})(1 - ERR_\zeta) \tag{10}$$

This is an heuristic method that assumes that $W_n$ will converge if we modify $\beta$ according to the gradient of $F_n$ and assume a linear relationship between $\beta$ and the equivalent stiffness $K$. Similarly, this method assumes a linear relationship between $\delta$ and the equivalent damping $C$. Consequently, $\delta$ is updated according to the gradient of $\zeta$ as well as according to the modification applied to $\beta$.

The resonance frequency of the axial joints was set to 8Hz. This value ensures that the joints are rigid enough to counter the inertial forces during fast swimming behaviors, with frequencies reaching up to 4.5 Hz [4, 5]. Indeed, lower values of resonance frequency would result in compliant joints and, conversely, in the inability to generate rapid and coordinated movements. On the other hand, higher values of resonance frequency would reduce the capability of external factors (i.e.; inertial, frictional and hydrodynamic forces) to counteract the torques generated by the muscles. In the limit case, the joint angles would only be the result of the muscle inputs, effectively approaching a position-controlled model. Overall, the choice of the resonance frequency was a trade-off between controllability of the model and responsiveness to external factors.

The damping factor of the axial joints was set to a value of 1.0, corresponding to the critically damped case. This specific damping value was chosen to avoid unwanted system responses; lower damping values led to uncontrolled motion in the caudal joints when the rostral ones were activated, resembling the dynamics of a chaotic double pendulum [6]. Conversely, higher damping values reduced the amplitude of the generated oscillations and slowed down the muscle dynamics. Similar damping factor values were also studied on a simplified single-joint model of the lamprey [1].

In Supp. Fig S1B, the convergence of the optimization procedure is illustrated. It can be observed that $ERR_{F_n}$ and $ERR_\zeta$ converge to 0 for all the axial joints. Similarly, Supp. Fig S1C shows the iterative refinement of the muscle parameters to improve the locomotor performance of the axial body. Interestingly, the muscle model parameters must differ widely across the different joints of the body for them to display the same target resonance frequency and damping factor.

This optimization highlights the interaction between muscle parameters and inertial properties, emphasizing the importance of precise parameter tuning to achieve the desired locomotor outputs. The findings also offer a framework for further research into the optimization of muscle parameters in biomechanical models, potentially leading to more refined simulations and a deeper understanding of locomotor dynamics in similar biological systems.

Notably, the delay between motoneurons activity and resulting kinematics output (i.e. the neuromuscular delay) in real salamanders displays a gradient along the axial body. Indeed, the neuromuscular delay increases in a rostro-caudal direction, displaying a discontinuity at the hip girdle [4, 7, 8]. This result, which was observed also in several fish species and the lamprey [9–11], suggests that the trunk and tail musculature could operate in different dynamic regimes. The latter could (in first approximation) display a higher damping factor which could justify the higher neuromuscular delay. Future studies may address the possibility to set different resonance frequencies and damping factors along the axial body in order to account for this phenomenon.
