## Supplementary material for "Balancing central control and sensory feedback produces adaptable and robust locomotor patterns in a spiking, neuromechanical model of the salamander spinal cord": S4_matching_swimming_kinematics

### Matching experimental swimming kinematics

In-vitro studies from *Pleurodeles Waltl* show that CPGs do not display any gradient in their properties along the trunk and tail networks [1–3]. Despite this observation, a distinct gradient of angle amplitudes is shown during anguilliform swimming [4–6]. Additionally, the musculature structure and the inertial properties change widely along the axial body of salamanders [7]. For these reasons, it is conceivable to assume that the overall force produced by muscle cells should also change along the body in order to account for the different mechanical properties and ranges of motion. In the model, this gradient is achieved by modulating the gain factor mediating the relationship between the membrane potential of the muscle cells and the input to the Ekeberg muscles ($G_{MC}$, see Eq 10 and Fig 2F in the main text).

The mechanical model was driven by sinusoidal signals provided as input to the Ekeberg muscles. The signals allowed to precisely control the frequency and phase lag of the muscle drives, matching the ones reported in [6] The gains were then optimized to align the amplitude of angle oscillations with observations made in-vivo [4–6].

With the optimized gains (i.e.; when matching the reference joint angles evolution), the obtained swimming behavior exhibited a notable similarity with the speed and lateral body displacements documented in *Pleurodeles Waltl* and other salamander species [4, 6]. In particular, the resulting lateral body displacements of the model fall within the ones reported in [6] (see Fig S1A). Similarly, the stride lengths obtained in simulation at different frequencies were comparable to the ones reported in [6]. Interestingly, salamanders show a wide level of variability in their swimming speed for a given frequency of movements [4]. Consequently, the stride lengths obtained in our model align well with the ones reported in real animals. The described matching of swimming metrics also ensures about the soundness of the drag coefficients employed in the hydrodynamic drag model (see Eq 16).

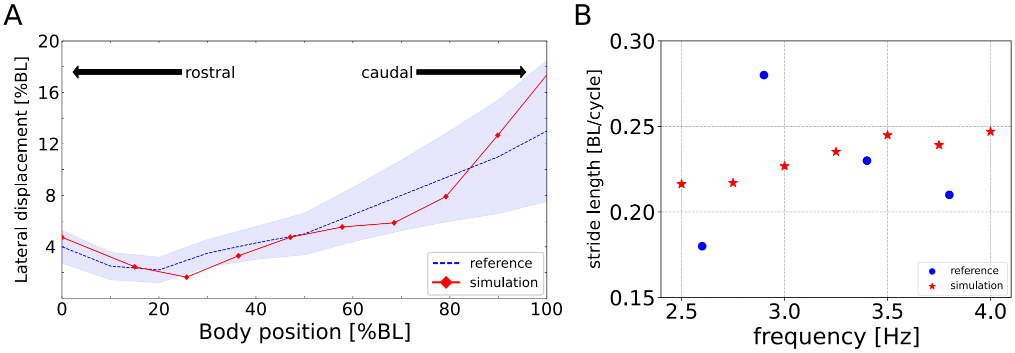


**Fig. S1. Optimization of mechanical model parameters. A**) shows the lateral displacements profile of the real and simulated salamanders. The blue line represents the average lateral displacements, expressed as percentage of body length (BL), observed along the spine in real salamanders. The blue area encompasses one standard deviation of difference from the average values. The red curve shows the corresponding lateral displacements obtained in the mechanical model when imposing the angles profiles recorded in [6] at a frequency of 3Hz. The obtained displacements closely resemble the experimental ones. **B**) displays the distribution of frequencies and stride lengths observed in real salamanders (blue dots) and in the current model (red stars) when imposing the angles profiles from [6] at different frequencies. Despite the simplicity of the drag hydrodynamic model, the speeds of the simulated salamander are close to the ones reported in real animals. These results validate the selected drag model parameters.
