## Supplementary material for "Balancing central control and sensory feedback produces adaptable and robust locomotor patterns in a spiking, neuromechanical model of the salamander spinal cord": S5_supplementary_results

#### Open loop swimming

##### Simulating fictive locomotion

Biological experiments typically by-pass the descending brain drive to directly investigate the capability of the spinal circuits to generate locomotor activity. Such experiments typically involve the use of N-methyl-D-aspartate (NMDA) baths to increase the excitability of isolated spinal cord sections [1, 2]. The generated activity is denoted "fictive locomotion". In order to simulate fictive locomotion, we carried out experiments where the axial central pattern generator (CPG) network was directly stimulated by an external current (see Fig S1). In each simulation, the CPG circuits where excited by an external current $I_{ext} = 20pA$, corresponding to the average rheobase current of the axial neurons. Different simulations where carried out to demonstrate the capability of the isolated network to produce oscillations at the level of the full cord, full hemicord (i.e.; one side of the network), isolated spinal segments and isolated spinal hemisegments.

In Fig S1A, the full spinal cord (i.e.; 40 equivalent segments) was stimulated. The corresponding activity resulted in a travelling wave of left-right alternating oscillations along the spinal cord, resembling the ones observed during fictive swimming in salamanders [1, 2]. Similarly, Fig S1B shows that the full hemicord can also independently produce oscillations. This property is a consequence of the recurrent ipsilateral excitation within the network, together with the adaptative nature of the CPG neurons. Again, this result is in accordance to fictive locomotion experiments in salamanders and lampreys [1–4].

In Fig S1C, the network was reduced from 40 segments to 2 equivalent segments (i.e.; from 4800 to 240 simulated neurons). The isolated spinal cord section was still able to produce left-right alternating oscillations. Additionally, the oscillations had a lower cycle duration compared to the full cord, consistently with [1]. Finally, in Fig S1D the network was reduced to 2 equivalent hemisegments. Again, the isolated spinal cord section was able to produce oscillations. Additionally, the hemisegmental oscillations had a lower rhythmicity, higher duty cycle and a higher frequency compared to the corresponding segmental activity. This is again in accordance with the experimental literature from fictive locomotion experiments in salamanders and lampreys [1, 5]

Notably, oscillations could still be seen in the evolution of the filtered spike count even when a single segment or hemisegment of the network was considered. In this case, however, the spiking activity was sparse and more difficult to interpret (not shown). Interestingly, [1] also reports very low periodicity (PTCC) values for isolated segments and hemisegments in the salamander. Similarly, [6] reports a direct proportionality between the number of considered segments and the quality of the corresponding activity in the lamprey, with very low PTCC values reported for a isolated ventral roots.

Overall, the simulations demonstrate the capability of the model to produce oscillations at the level of full cord, full hemicord, isolated segments and isolated hemisegments. The results align well with the available literature from fictive locomotion experiments in salamanders and lampreys.

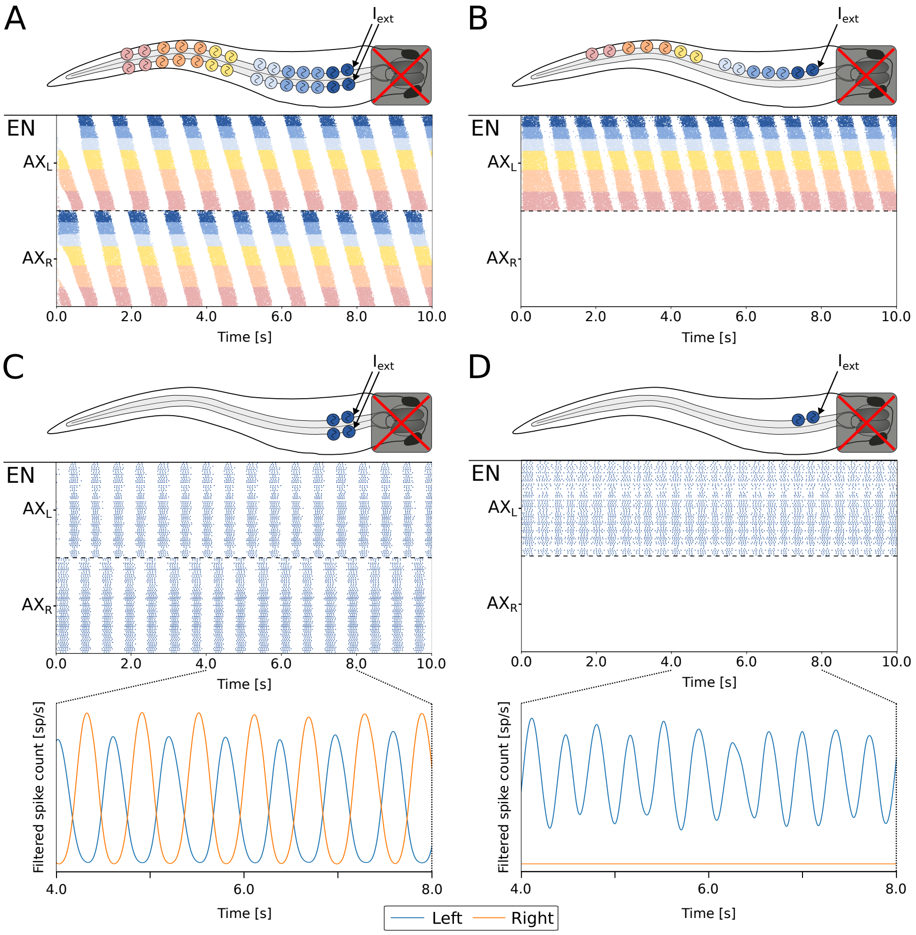


**Fig. S1. Simulation of fictive locomotion.** Fictive locomotion was obtained in the model by directly stimulating the central pattern generator network (black arrows) and by-passing the reticulospinal neurons (red cross). The external current ($I_{ext}$)) was set to 20pA. Raster plots represent the spike times of excitatory (EN) neurons of the axial (AX) network. The upper half represents the neurons on the left side (L), the lower half represents the neurons on the right side (R). In C and D, the bottom figure shows the filtered spike count for the left (blue) and right (orange) side of the network. **A**) shows the pattern obtained when activating the full spinal cord (i.e.; 40 segments). **B**) shows the pattern obtained when activating a full spinal hemicord (i.e.; 40 hemisegments). **C**) shows the pattern obtained when activating two isolated spinal segments. **D**) shows the pattern obtained when activating two isolated spinal hemisegments.

#### Effect of descending drive

The response of the open-loop network was studied across varying stimulation values targeting the axial reticulospinal neurons (RS). The stimulation amplitude was varied in the range $[4.0, 8.0] pA$ for 20 different instances of the network, built with different seeds for the random number generation. The results are reported in terms of mean and standard deviation of the corresponding computed metrics.

The analysis revealed a triphasic response of the network within the range of considered stimulation amplitudes (see Fig S2A). Notably, when the stimulation amplitude was lower than the rheobase current of the RS neurons (equal to $4.0pA$ on average), the CPG network did not receive any descending drive and thus remained silent. For stimulation values close to the rheobase current, the network displayed aperiodic patterns and disorganized oscillations. This resulted in low periodicity values (PTCC, see Eq 4 in S1 File), indicating diminished network rhythmicity (see Fig S2B, top panel). For intermediate stimulation values, the network manifested traveling waves of coordinated activity alongside a distinct left-right alternation (see Fig S2B, middle panel), analogous to the motor patterns observed during swimming [7]. This phase represents a functional regime wherein the network accurately replicates rhythmic motor outputs associated with swimming. At higher stimulation values, the network deviated from oscillatory behavior, with both hemisegments exhibiting sustained tonic activity (see Fig S2B, bottom panel). In this phase, the combined action of commissural inhibition and neuronal adaptation was insufficient to counteract the persistent excitation induced by the elevated background drive, resulting in a loss of rhythmic oscillations across the hemisegments. Note that, in this regime, the network loses the capability to produce stable rhythms (i.e.; $PTCC \approx 0$) and the corresponding frequency ($f_{neur}$) and total wave lag ($TWL$) metrics are therefore unreliable.

Within the oscillatory regime delineated by intermediate stimulation levels, a linear relationship emerged between the stimulation drive and the frequency of oscillations (see Fig S2A, middle panel). Additionally, the frequency of oscillations varied in the range $[1.0, 4.0]Hz$, aligning with empirical findings from salamander studies involving MLR stimulation [8]. Conversely, the total wave lag ($TWL$, see Eq 3 in S1 File), initially increased with rising drive, plateauing around a value of 1.2 (see Fig S2A, bottom panel). This $TWL$ value was in the lower range of those reported for the swimming kinematics in the *Ambystoma Mexicanum* (ranging from 1.2 to 2.1 in [9]). Conversely, the $TWL$ for the epaxial EMG activity during swimming in *Pleurodeles waltl* was markedly lower (0.5 in [10], 0.74 in [2]). Notably, these experiments were carried out on behaving animals naturally receiving sensory feedback during locomotion. Interestingly, isolated *Pleurodeles waltl* spinal cord preparations were shown to display much higher $TWL$ values when stimulated with NMDA baths (up to 2.5 in [1]). As shown in the section "Closed loop swimming" in the main text, the introduction of sensory feedback can lower the $TWL$ values to match the experimental findings. This result also provides a possible explanation for the mismatch between isolated and intact spinal cord recordings.

The observed drive-frequency relationship parallels the findings from prior simulation studies conducted at higher abstraction levels, with coupled nonlinear oscillator models [11]. This consistency strongly suggests that for the salamander the assumptions of models at higher levels of abstraction are well justified through a chain of models, ranging from detailed bio-physical models (e.g., [12]) through formal spiking models (e.g., [13] and present model), all the way to more abstract controllers [11, 14]. Unless stated otherwise, all the analyses in this work impose a stimulation amplitude of 5.7pA to the axial RS neurons. With the selected drive, the network reliably generates stable oscillations with an average frequency of $3.1Hz$ and a total wave lag of $1.27BL/cycle$.

#### Effect of external noise

The swimming network was simulated with different noise levels to investigate its robustness (see Fig S3A). The noise amplitude varied in the range $[0.0, 6.25]pA$ and each simulation was carried out for 20 instances of the network, built with different seeds for the random number generation. It was found that the axial CPG network could tolerate high levels of noise without notably altering the resulting pattern (see

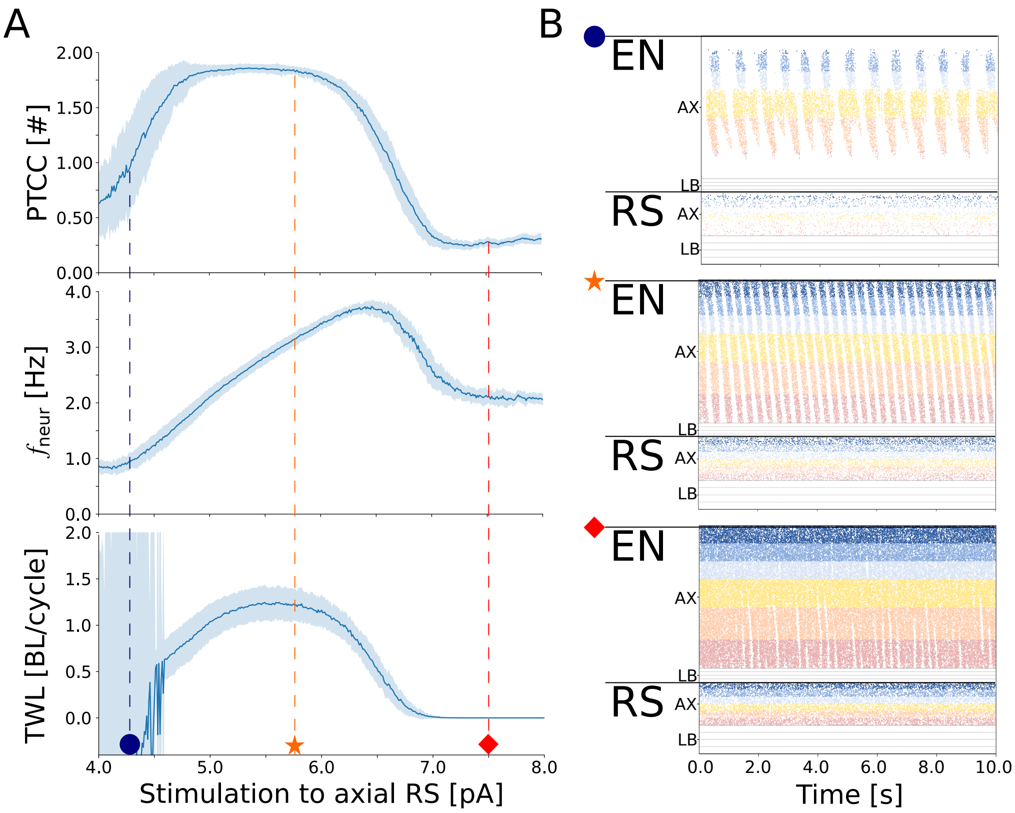


**Fig. S2. The triphasic response to the stimulation amplitude to the axial RS neurons. A**) displays the effect of the stimulation amplitude provided to the axial RS neurons, on the open loop swimming network. The stimulation was varied in the range $[4.0, 8.0]pA$. From top to bottom, the plots show the relationship between the level of stimulation and the periodicity (PTCC), frequency ($f_{neur}$) and total wave lag ($TWL$) of the neural signals, respectively. In each plot, the thick line represents the average metric computed across 20 different instances of the network, built with different seeds for the random number generator. The shaded area denotes one standard deviation of distance from the mean. Note that, for $PTCC \approx 0$, the network loses its capability to produce stable rhythms and the values of $f_{neur}$ and $TWL$ are therefore unreliable. **B**) shows the response of the network for three different drive amplitudes to the axial RS neurons (equal to $4.25pA$, $5.75pA$ and $7.5pA$, respectively). The raster plots represent the spike times of excitatory neurons (EN) and reticulospinal neurons (RS). For the axial network (AX), only the activity of the left side is displayed. For the limb network (LB), only the activity of the flexor side is displayed. For low stimulation values (blue dot), the network displays unpatterned activations. For intermediate stimulation values (orange star), the network displays travelling waves of left-right activity. For high stimulation values (red square), the network displays tonic activity.

Fig S3B). Indeed, despite a high noise level of $4.5pA$, the network could generate a coordinated swimming pattern that resembled the one without external disturbance from Fig 3A1,A2 in the main text.

This robustness is likely due to the diversity in the neuronal parameters of the CPG populations, as suggested by [15]. The diverse neuronal parameters may provide a form of network heterogeneity that helps buffer against the disruptions caused by noise, ensuring functional outputs are maintained under various conditions. However, there is a limit to this tolerance. When noise levels were exceedingly high (i.e., greater than 5pA), a noticeable decline in the rhythmic pattern was observed, leading to a cessation of oscillatory activity and, consequently, low values of PTCC (see Fig S3A).

The findings from this simulation highlight the axial CPG network's robust nature in handling noise and also outline the boundaries of this robustness.

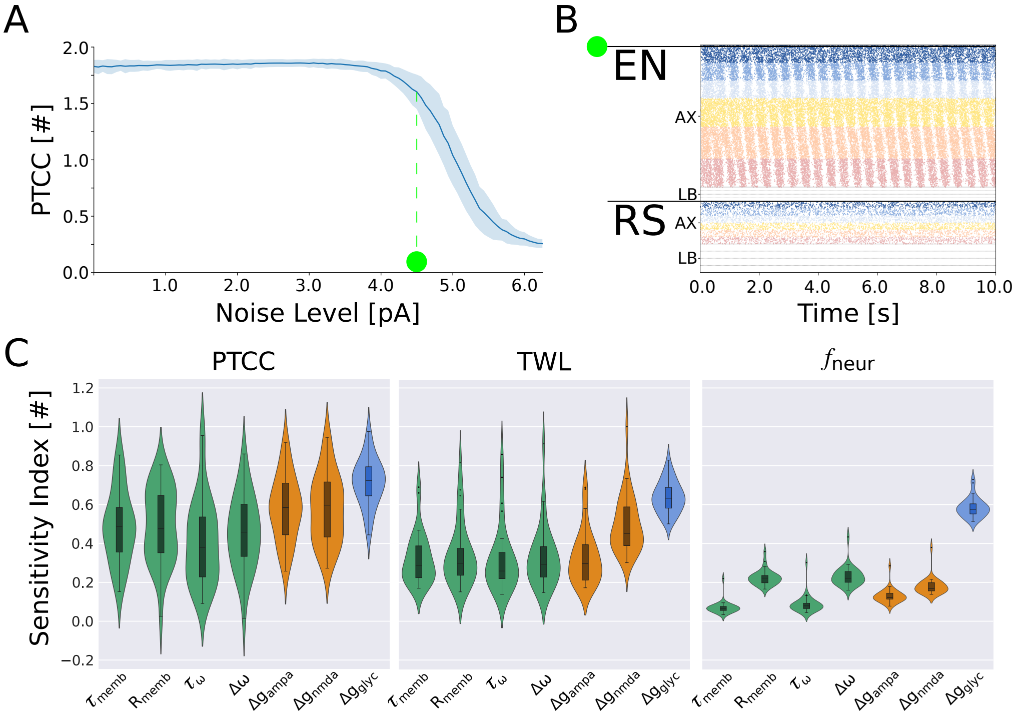


**Fig. S3. The network's inherent robustness to noise and sensitivity analysis. A**) displays the response of the swimming network to different values of noise level, which was varied in the range $[0.0, 6.25]pA$. The thick line indicates the average PTCC value over 20 instances of the network, built with different seeds for the random number generator. The shaded area around the curve indicates the region within one standard deviation of distance from the mean. **B**) shows the response of the network for a noise level of 4.5pA. The raster plots represent the spike times of excitatory neurons (EN) and reticulospinal neurons (RS). For the axial network (AX), only the activity of the left side is displayed. For the limb network (LB), only the activity of the flexor side is displayed. **C**) shows the results of the sensitivity analysis performed on the open loop network. The x-axis lists the neuronal and synaptic parameters changed during the analysis. The y-axis shows the distribution of the corresponding total-order sensitivity indices for 30 different instances of the network, built with different seeds for the random number generator. The three plots show the sensitivity indices with respect to the periodicity (PTCC), total wave lag (TWL) and frequency ($f_{neur}$), respectively. It can be noted that the weight of inhibitory connections $\Delta g_{glyc}$ is the parameter with the highest influence on the considered metrics.

#### Open loop sensitivity analysis

The sensitivity analysis pointed out the important role of commissural inhibitory connections in modulating rhythmicity (PTCC), total wave lag (TWL), and frequency ($f_{neur}$) of the network's locomotor output (see Fig S3C). The figure reports the distribution of the total order sensitivity indices computed across 20 instances of the network, built with different seeds for the random number generation. The indices were computed for all the neuronal and synaptic parameters with respect to the periodicity, total wave lag and frequency of the neural activity. Indeed, for all the considered metrics, the synaptic weight of inhibitory connections ($\Delta g_{glyc}$) resulted in the highest total-order sensitivity index among all neuronal and synaptic parameters. The result was particularly evident for $f_{neur}$, where $\Delta g_{glyc}$ showed a very strong modulatory effect. The nature of the modulation of $f_{neur}$, $TWL$ and $PTCC$, in terms of increase or decrease of the corresponding metrics, is discussed in detail in the next section.

This analysis sheds light on the critical role of commissural inhibitory connections as major modulators of network rhythmicity and locomotor tempo. The results support the escape-from-inhibition hypothesis [16, 17], where rhythmic locomotor patterns arise from a balance of excitatory and inhibitory interactions within the network.

#### Effect of commissural inhibition

In the previous section, we discussed the critical role of the CPG commissural inhibition in generating patterned locomotor activity. It notably affects the frequency, phase lag, and periodicity of the network, making it crucial to understand the range of inhibition strengths that allow rhythm generation. We conducted an analysis by varying the strength of commissural inhibition between hemisegments and studying the corresponding periodicity (PTCC), frequency ($f_{neur}$) and total wave lag (TWL) values displayed by the network (see Fig S4A). The commissural inhibition strength weight was varied in the range $[0.0, 4.0]$. The stimulation amplitude to axial RS neurons was set to 5.7pA. The curves display the average PTCC (in blue), $f_{neur}$ (in red) and TWL (in green) values across 20 instances of the network, built with different seeds for the random number generator. The shaded areas highlight the metrics values within one standard deviation from the average. The figure reveals three distinct regions based on inhibition strength (see Fig S4B). At low inhibition strength, the descending drive overpowers the commissural inhibition, leading to bilateral tonic activation in the network (see Fig S4B1). At high inhibition strength, neuronal adaptation is not enough to allow the contralateral side to escape from the inhibition, resulting in a 'winner takes all' configuration where one side becomes tonically active while the opposite side remains silent (see Fig S4B3). In this configuration the network is effectively bi-stable and can converge to either the left-side-winning or right-side-winning configuration depending on the initial conditions of the neurons. At intermediate inhibition strength, the network exhibits stable oscillations (i.e. $PTCC > 1.0$), as indicated by the plateau in the value of PTCC (see Fig S4B2). Interestingly, in the oscillatory regime the frequency of the oscillations decreased monotonically with increasing values of inhibition stregth. Conversely, the TWL showed an initial increase up to $TWL \approx 1.3 BL/cycle$ followed by a sharp decrease to $TWL \approx 0.0 BL/cycle$ (i.e. a standing wave of activity).

Notably, when the commissural inhibition weight is set to 0, the model becomes equivalent to two independent hemicord networks, each activated by descending RS drive. In this configuration, hemicords would be expected to generate independent oscillations, similarly to what observed when simulating hemicords activity during fictive locomotion (see section "Simulating fictive locomotion" and Fig S1B). Conversely, the network in Fig S4B1 did not produce oscillations in this configuration. Notably, there is an active debate in the lamprey literature regarding the capability of the brain-activated network to produce hemicord oscillations [18–20]. Indeed, [19] showed that eliminating left-right reciprocal coupling could abolish the sensory-induced (and brain-driven) generation of locomotor activity, which was substituted by a tonic network activation. Interestingly, a possible explanation of this mismatch between sensory-induced and bath-induced activity could reside in a high RS activation (and therefore descending drive) caused by the sensory stimulation performed in [19]. Indeed, hemicord oscillations can be obtained in the current model also in the RS-activated network when decreasing the external stimulation level to 4.75pA (see Fig S5).

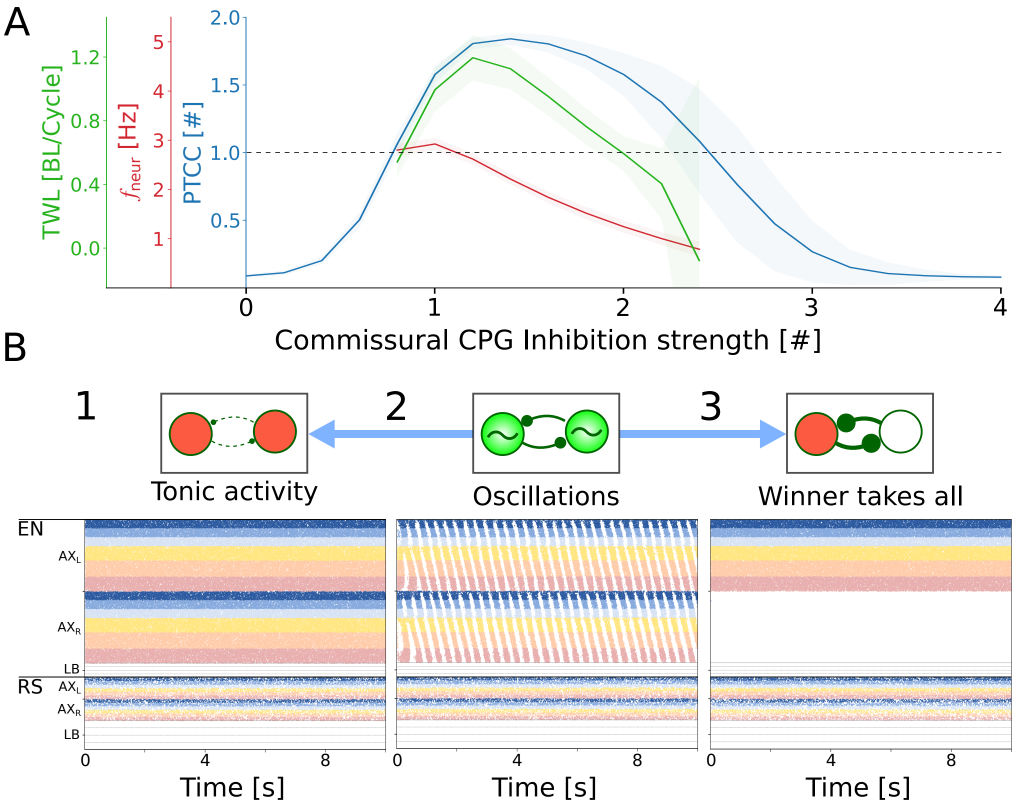


**Fig. S4. Commissural inhibition governs rhythm generation, frequency and intersegmental phase lag. A**) shows the dependency of periodicity (PTCC, in blue), frequency ($f_{neur}$, in red) and total wave lag (TWL, in green) on the strength of the CPG commissural inhibition. The inhibition strength was varied in the range $[0.0, 4.0]$. The external RS drive to the axial network was kept constant and equal to 5.7pA. The thick lines represent the mean across 20 instances of the network, built with different seeds for the random number generator. The shaded areas represent one standard deviation of difference from the mean. The values of $f_{neur}$ and TWL are shown only for combinations where the average PTCC was greater than 1.0 (dotted line). **B**) shows the three regimes displayed by the network depending on the level of commissural inhibition strength. Raster plots represent the spike times of excitatory (EN) and reticulospinal (RS) neurons of the axial (AX) and limbs (LB) sub-networks. For each population, the upper half represents the neurons on the left side (L), the lower half represents the neurons on the right side (R). For low inhibition values (B1) both sides of the network are tonically active. For intermediate inhibition values (B2) the network displays rhythmic left-right alternating waves of activation. For high inhibition values (B3) one side of the network is tonically active causing the other one to remain silent (”winner takes all” configuration).

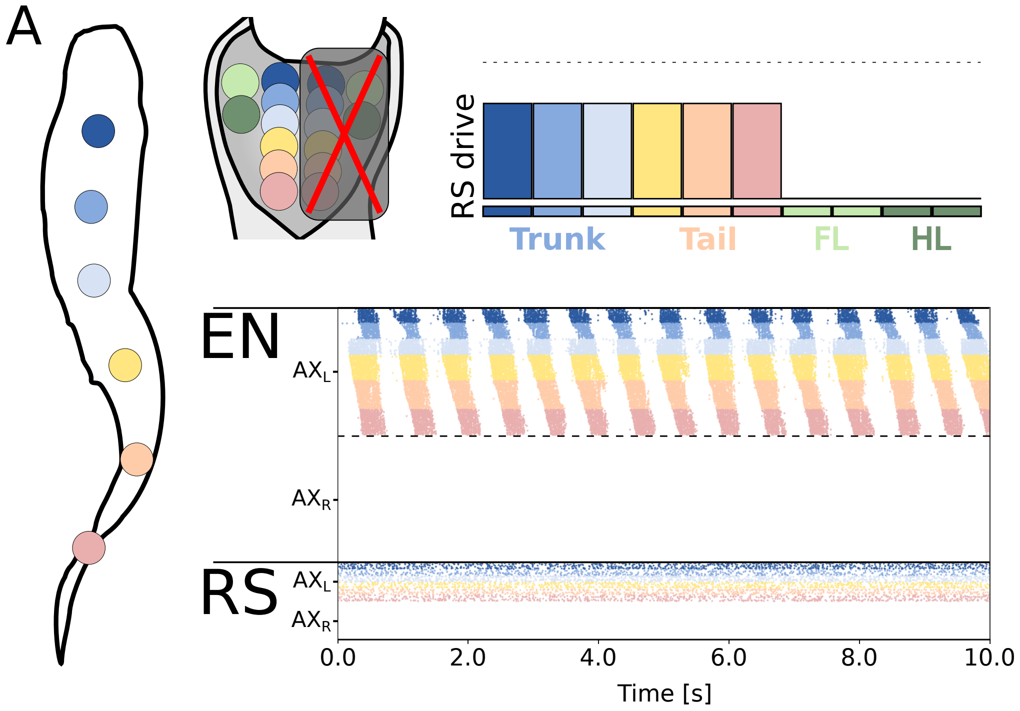


**Fig. S5. Hemicord oscillations induced by descending RS drive. A**) shows the pattern emerging from a stimulation of left axial reticulospinal (RS) neurons. The right RS neurons are kept silent (red cross). The level of stimulation provided to the 6 axial RS modules was equal to 4.75pA. The dotted line indicates a value of 5.7pA, used by most simulations in this work. The raster plot represents the spike times of excitatory neurons (EN) and reticulospinal neurons (RS) of the axial network (AX). For each population, the upper half represents the neurons on the left side (L), the lower half represents the neurons on the right side (R). The hemicord network can generate stable oscillations with descending RS drive, provided that the external RS stimulation is sufficiently low.

### Closed loop swimming

#### Closed loop sensitivity analysis

The sensitivity analysis was repeated in closed loop control of the mechanical model with different values of $\omega_{PS}$ (see Fig S6). The varied parameters included the same neuronal parameters studied for the open loop network. Additionally, the value of $\omega_{PS}$ was also modified by the analysis around its nominal value. The studied nominal values for $\omega_{PS}$ ranged from 0 (no sensory feedback case) to 1.5 (high sensory feedback case). A separate sensitivity analysis was carried out for each nominal $\omega_{PS}$ value. Each panel of Fig S6 shows the distribution of the total-order sensitivity indices computed for the different neuronal and mechanical parameters with respect to the correspondent metric. The distributions were obtained from 20 instances of the network built with different seeds for the random number generation.

Studying the effect on the network's periodicity (PTCC), it can be seen that increasing $\omega_{PS}$ led to a decrease of the total-order sensitivity indices for all the remaining parameters. This result could indicate that the network becomes less dependent on a specific parameter tuning to generate stable oscillations. On the other hand, for $\omega_{PS} > 0.5$, the sensitivity index for the feedback weight itself tends to decrease. Indeed, for high $\omega_{PS}$ values, the effect of sensory feedback reaches a saturation and the network is not affected by small variations around its nominal value. Interestingly, the commissural inhibition weight ($\Delta g_{glyc}$) was consistently the parameter with the highest total-order sensitivity index. This indicates that, regardless of $\omega_{PS}$, the network's periodicity is still primarily affected by changes in the inhibition strength.

The described trends are even more evident for the sensitivity indices computed with respect to the total wave lag (TWL, see Eq 3 in S1 File). Also in this case we observe a large decrease in the sensitivity indices of all the parameters with increasing values of $\omega_{PS}$. Similarly to the previous case, the sensitivity index for the feedback weight reaches a maximum for $\omega_{PS} = 0.5$ and then drops for higher values. Indeed, higher values of $\omega_{PS}$ lead to a quasi-standing wave of axial activation (see Fig 7C in the main text) independently of small variation of the feedback weight value.

Finally, similar observations can be made for the sensitivity indices of the studied parameters with respect to the frequency ($f_{neur}$), forward speed ($V_{fwd}$) and cost of transport (COT) of the generated activity. Overall, it was shown that intermediate values of $\omega_{PS}$ reduce the dependency of the locomotor circuits on a precise selection of the neuronal and synaptic parameters. On the other hand, excessive values of sensory feedback strength dominate the network activity and reduce the capability to actively control its behavior.

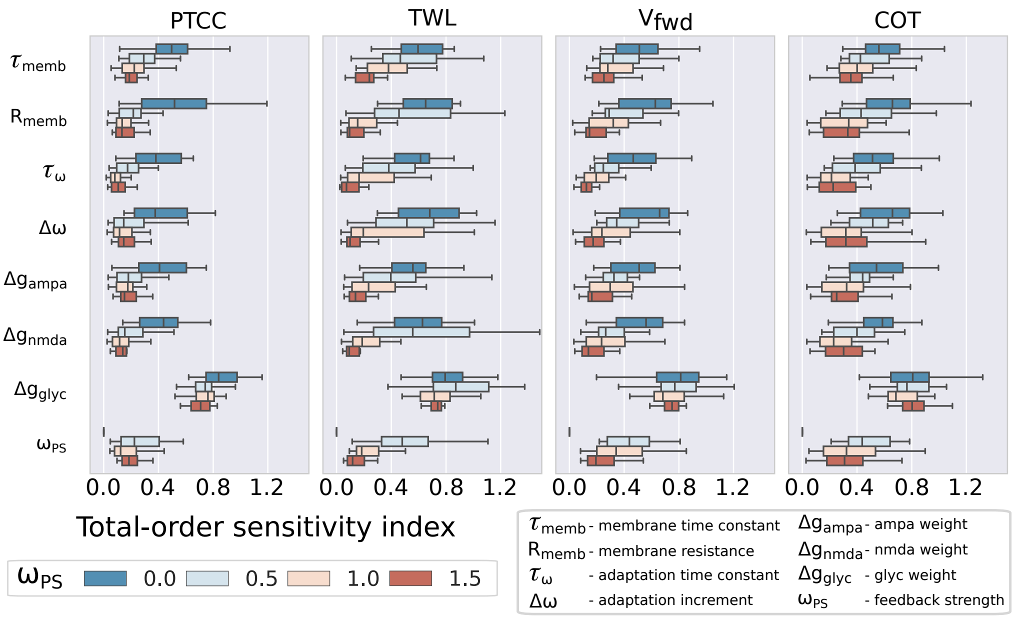


**Fig. S6. Sensory feedback reduces the sensitivity to neuronal and synaptic parameters.** Each panel displays the distribution of the total-order sensitivity indices computed for the different network parameters (listed on the y-axis and legend) with respect to the periodicity (PTCC), total wave lag (TWL), forward velocity ($V_{fwd}$) and cost of transport (COT). The colors are associated to different values of $\omega_{PS}$, reported in the legend.

#### Supplementary figures

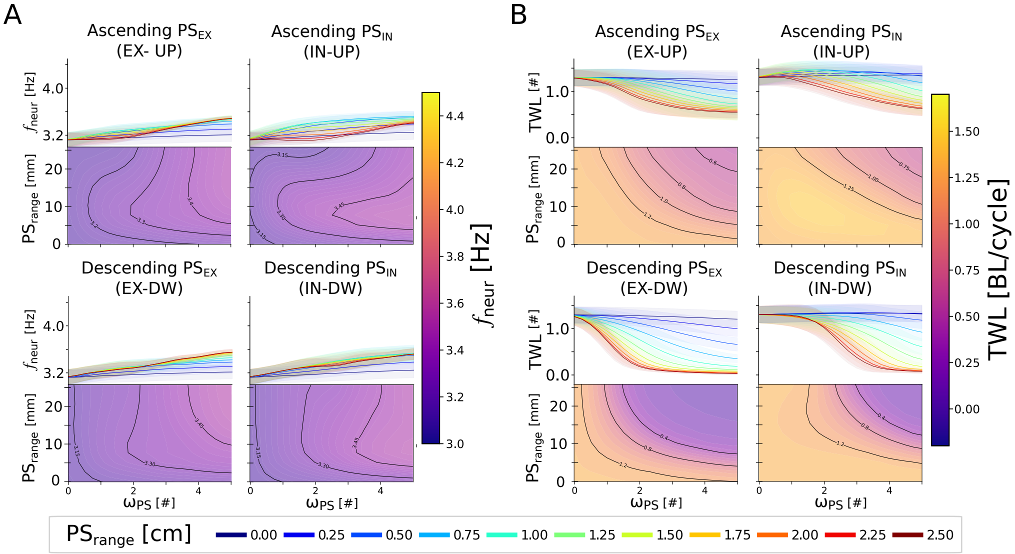


**Fig. S7. Effect of feedback topology on the closed loop swimming network for $\theta_{RH}^{\%} = 50\%$.** Panels A and B show the effect of different PS topology patterns and weights on the frequency ($f_{neur}$) and total wave lag ($TWL$) of the network's activity, respectively. The studied topologies include ascending excitation (EX-UP), descending excitation (EX-DW), ascending inhibition (IN-UP) and descending inhibition (IN-DW). The range of the connections (PS range) was varied in the range $[0, 2.5]mm$ (i.e., between 0 and 10 equivalent segments). Similarly, the PS synaptic weight ($\omega_{PS}$) was varied in the range $[0, 5]$. **A**) shows the modulation of the oscillations' frequency ($f_{neur}$). In each quadrant, the contour plot shows the dependency of the $f_{neur}$ on PS range and $\omega_{PS}$. On top, the projection of the relationship in the $f_{neur} - \omega_{PS}$ plane is displayed to better highlight the asymptotical values reached by $f_{neur}$. The different curves represent the frequency expressed by the network with different PS connection ranges (i.e.; horizontal cuts of the contour plot), whose value is reported in the legend. The thick lines represent the mean metric values across 20 instances of the network, built with different seeds for the random number generator. The shaded areas represent one standard deviation of difference from the mean. **B**) follows the same organization as the previous panel to represent the effect of feedback topology on the neural total wave lag ($TWL$).

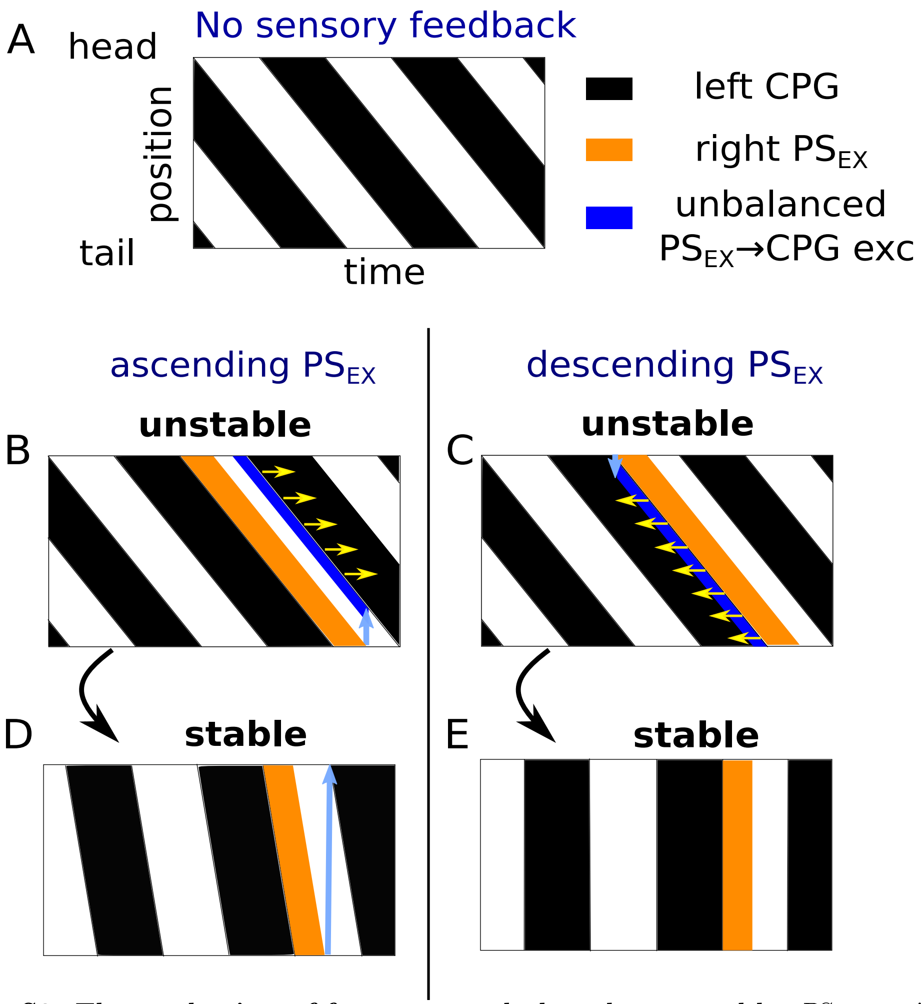


**Fig. S8. The mechanism of frequency and phase lag control by $PS_{EX}$ . A-E)** shows the stereotypical CPG activities during swimming in a brief time window (black=active/firing, white=inactive) sorted according to the CPG positions (top=head positions, bottom=tail positions). **A** shows the activities of CPGs with no feedback (in open loop). **B-E** also shows the activities of CPGs and right $PS_{EX}$s (orange) in closed loop. B-D and C-E show subsequent time-windows at starting and final stages of activities from the time when $PS_{EX}$ is turned on, respectively. The light blue arrows represent the $PS_{EX}$ projections in the ascending (B-D) and descending (C-E) directions to the closest CPG neurons on the ipsilateral sides. These indicate the end/start of a region of unbalanced excitation from $PS_{EX}$ to the CPGs (dark blue areas). This imbalance causes a shift in the left-right activity transition of the corresponding CPGs according to the direction indicated by the yellow arrows. According to our mechanistic explanation, the system evolves to minimize the unbalanced $PS_{EX}$ excitation to the CPGs (i.e., the dark blue area). In B ascending $PS_{EX}$ excitation is higher for rostral CPGs. This causes these neurons to delay their inactivation, which in turn delays the activation of the rostral left CPG neurons, causing a global decrease in the network wave lag, analogously to the $PS_{IN}$ case (main text). This decrease in the wave lag will continue until the wave reaches a stable state with positive wave lag (D). In C the $PS_{EX}$ excitation is higher for caudal CPGs. This unbalance causes caudal CPGs to anticipate their activation, which decreases the overall network wave lag. In this case a stable oscillatory traveling wave is only guaranteed when the wave lag is approximately zero.

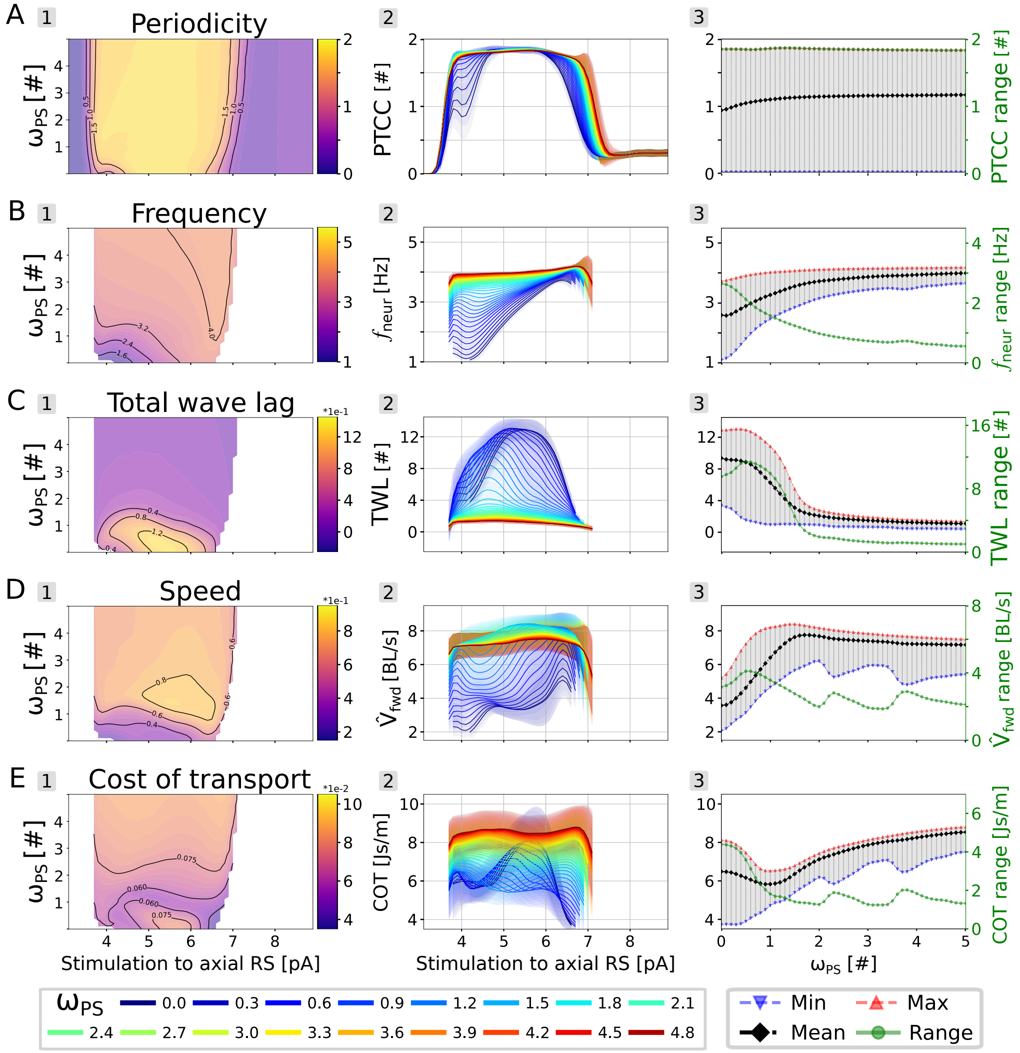


**Fig. S9. Effect of drive on the closed loop swimming network for $\theta_{RH}^{\%} = 50\%$. A**) shows the effect of the stimulation amplitude and feedback weight ($\omega_{PS}$) on the periodicity (PTCC) of the generated activity. The stimulation amplitude was varied in the range $[3, 9]pA$. The feedback weight was varied in the range $[0, 5]$. The displayed data is averaged across 20 instances of the network, built with different seeds for the random number generator. In A2, the curves differ by their corresponding value of $\omega_{PS}$, reported in the bottom legend. The shaded areas represent one standard deviation of difference from the mean. A3 shows the minimum (in blue), maximum (in red) average (in black) and range (in green) for the PTCC values across all the considered stimulation values. The range values are reported on the y-axis on the right. Panels (B-E) follow the same organization to display the effect on the oscillation frequency ($f_{neur}$), total wave lag (TWL), forward velocity ($V_{fwd}$) and cost of transport (COT). Combinations leading to PTCC values lower than 1 were not displayed.

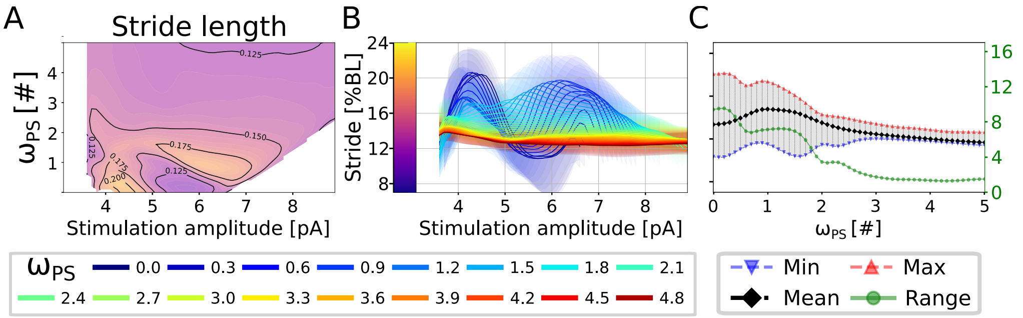


**Fig. S10. Effect of drive on the stride length of the closed loop swimming network. A**) shows the effect of the stimulation amplitude and feedback weight ($\omega_{PS}$) on the stride length (Stride) of the generated activity. The stimulation amplitude was varied in the range $[3, 9]pA$. The feedback weight was varied in the range $[0, 5]$. The rheobase angle ratio was set to $\theta_{RH}^{\%} = 10\%$. The displayed data is averaged across 20 instances of the network, built with different seeds for the random number generator. In **B**), the curves differ by their corresponding value of $\omega_{PS}$, reported in the bottom legend. The shaded areas represent one standard deviation of difference from the mean. **C**) shows the minimum (in blue), maximum (in red) average (in black) and range (in green) for the stride length values across all the considered stimulation values. The range values are reported on the y-axis on the right.

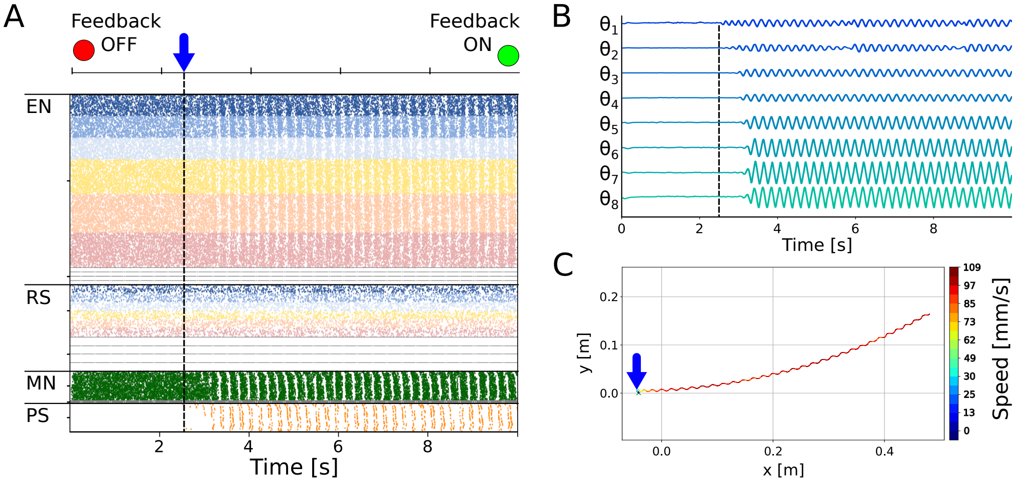


**Fig. S11. Restoring rhythmic activity with high levels of noise. A**) shows the switch effect with a noise level of 8.75pA. The raster plot represents the spike times of excitatory neurons (EN), reticulospinal neurons (RS), motoneurons (MN) and proprioceptive sensory neurons (PS). Only the activity of the left side of the network is displayed. The rheobase angle ratio was set to $\theta_{RH}^{\%} = 10\%$. At time=2.5s (blue arrow), the sensory feedback weight is changed from $\omega_{PS} = 0$ (Feedback OFF) to $\omega_{PS} = 2.0$ (Feedback ON), restoring a left-right alternating rhythmic network activity. **B**) shows the axial joint angles during the simulation in A. Each angle is shown in the range [-25°, +25°]. **C**) shows the center of mass trajectory during the simulation in A. Activating feedback restored forward swimming.

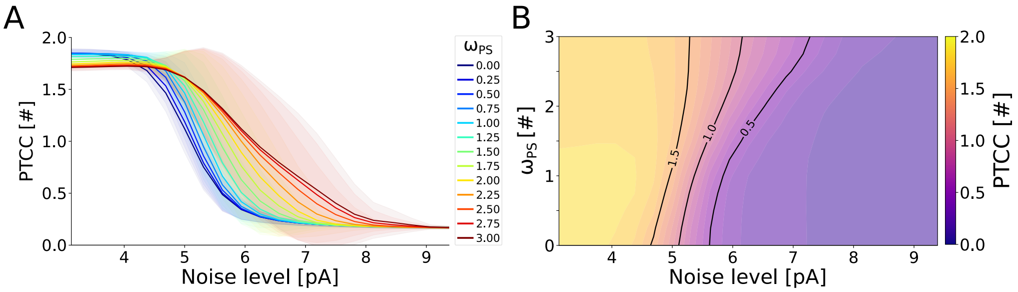


**Fig. S12. Effect of noise on the closed loop swimming network for $\theta^{\%}_{RH} = 30\%$. A**) shows the effect of noise amplitude on the rhythmicity of the network (PTCC) with different levels of sensory feedback weight ($\omega_{PS}$). The noise level was varied in the range $[3.0, 9.5]pA$. The feedback weight was varied in the range $[0, 3]$. The different curves represent the average PTCC values across 20 instances of the network (built with different seeds for the random number generator) for different levels of $\omega_{PS}$ (shown in the legend). The shaded areas represent one standard deviation of difference from the mean. **B**) displays the noise-feedback relationship in 2D.

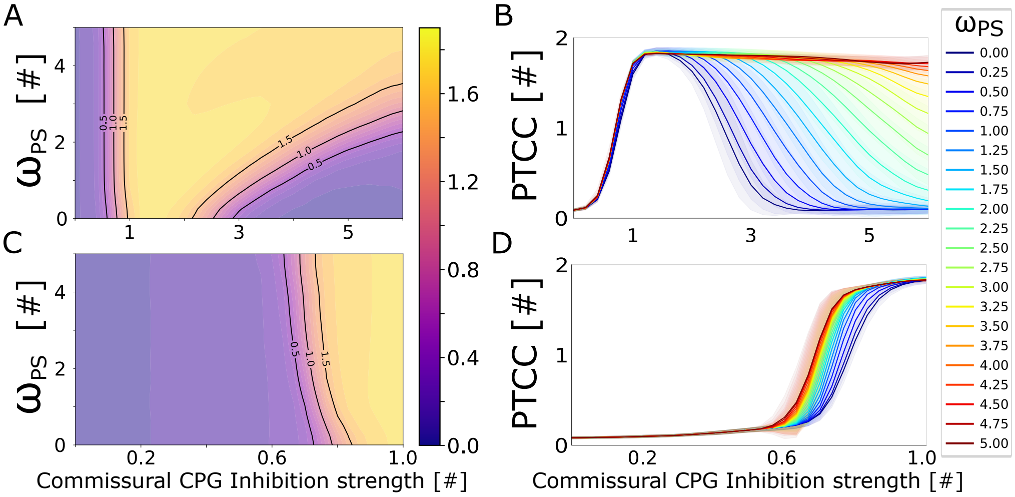


**Fig. S13. Effect of CPG commissural inhibition on the closed loop network for $\theta^{\%}_{RH} = 50\%$. A**) shows the dependency of the periodicity (PTCC) on the inhibition strength and on the feedback weight ($\omega_{PS}$). The commissural inhibition strength was varied in the range $[0, 6]$. The feedback weight was varied in the range $[0, 5]$. The displayed values are averaged across 20 instances of the network, built with different seeds for the random number generator. **B**) shows the inhibition-feedback relationship in 1D. The different curves represent the average PTCC values for different levels of $\omega_{PS}$ (shown in the legend). The shaded areas represent one standard deviation of difference from the mean. Panels C and D follow the same organization as the previous panels to highlight the inhibition-feedback relationship for $\omega_{PS}$ values in the range $[0, 6]$. **E**) displays the distribution of the total-order sensitivity indices computed for the different network parameters (listed on the y-axis) with respect to the PTCC, total wave lag (TWL), forward velocity ($V_{fwd}$) and cost of transport (COT). The colors are associated to different values of $\omega_{PS}$, reported in the legend.
